# DISC1 regulates astrogenesis in the embryonic brain via modulation of RAS/MEK/ERK signaling through RASSF7

**DOI:** 10.1101/032243

**Authors:** Shukun Wang, Qingli Liang, Huimin Qiao, Hong Li, Tianjin Shen, Fen Ji, Jianwei Jiao

## Abstract

Disrupted in Schizophrenia1 (*DISC1*) is known as a high susceptibility gene of schizophrenia. More recent studies have connected schizophrenia with glia defects and dysfunction. However, it is unclear whether there is connection between DISC1 and gliogenesis defect. Thus, a precise understanding of DISC1 (a ubiquitously expressed brain protein) on astrogenesis in the late stage of embryonic mouse brain development needs to be deeply investigated. Here, we show that suppression of DISC1 expression represses astrogenesis *in vitro* and *in vivo*, whereas DISC1 overexpression substantially enhances the process. Furthermore, mouse and human DISC1 overexpression rescued astrogenesis defects caused by DISC1 knowndown. Mechanistically, DISC1 activates downstream RAS/MEK/ERK signaling pathway via directly associating with the C terminal domain of RASSF7, a RAS association protein. Also, the pERK complex undergoes nuclear translocation and influences the expression of genes related to astrogenesis. Briefly, our results demonstrate that DISC1 regulates astrogenesis by modulating RAS/MEK/ERK signaling via RASSF7 and provide a framework for understanding how DISC1 dysfunction leads to neuropsychiatric diseases.

## INTRODUCTION

Disrupted-in-Schizophrenia 1 (*DISC1*) was originally identified in chromosome translocation (1; 11) (q42.1; q14.3) in a Scottish family with a high susceptibility to schizophrenia and several other psychiatric disorders (Hayashi-Takagi et al., 2010; Millar et al., 2000). It is a crucial element of the microtubule-associated dynein motor complex maintaining normal microtubular dynamics (Kamiya et al., 2005). The C-terminal mutant DISC1 or expression of mutant DISC1 in mouse brains lead to the manifestation of schizophrenia-like behavioral phenotypes (Li et al., 2007; Pletnikov et al., 2008).

Previous studies have revealed that DISC1 is important in embryo and adult brain development. Kamiya et al discovered that DISC1 depletion impairs neurite outgrowth (Kamiya et al., 2005). Enomoto *et al*. reported that DISC1 regulates axonal development via interacting with protein girdin (Enomoto et al., 2009). Mao *et al*. provided that DISC1 deficiency decreases the progenitor pool and results in premature neuronal differentiation via regulating GSK3β/β-catenin signaling. DISC1 dysfuntion in the dentate gyrus of adult brain suppresses neural progenitor proliferation and causes hyperactive and depressive behaviors (Mao et al., 2009a). Kim *et al*. pointed that DISC1 modulates AKT-mTOR signaling and then regulates neuronal development (Kim et al., 2009). Most researches of DISC1 are focused on the effects on neuron, but few studies are paid attention to the regulation of DISC1 on astrocyte.

Astrocytes, the most abundant type of glial cells in the nervous system, impact the function of surrounding neurons in various means (Lopez-Hidalgo and Schummers, 2014). During forebrain development, neural stem cells generate neuron first, followed by astrocyte, and finally oligodendrocytes. This temporal sequence of neocortex development is tightly regulated by both intrinsic programs and extracellular signals (Brambilla et al., 2013). Astrocytes are originated from progenitor cells in the SVZ at the late embryonic days and reach the peak level at early postnatal life (Sauvageot, 2002). Glial progenitors, derived from common progenitors, can differentiate into astrocytes (Huse and Holland, 2010).

Astrocytes physically interacts with neurons, offer metabolic support, buffers the extracellular ions, and releases trophic factors and neurotransmitters after modulating information procession and signal transmission (Annunziato et al., 2013; Barros, 2013; Lopez-Hidalgo and Schummers, 2014). It is reported that dysfunctional astrocytes affect neighboring neurons and then result in neurodegeneration and brain disease (Avila-Munoz and Arias, 2014). Moreover, it is revealed that the disruption of astrocyte was found in schizophrenic patients by observing vast increase of extracellular matrix proteins in the brain (Pantazopoulos et al., 2010). And several depressive mouse models show defects in glial cell density (Molofsky et al., 2012). Therefore, we hypothesize that DISC1 dysfunction maybe affect astrogenesis and then result in brain defect. Therefore, it is important to understand the molecular mechanism by which DISC1 regulates astrogenesis.

In neurogenesis, KIAA1212 is proved to associate with both CT and MD domains of DISC1 when DISC1 regulates new neuron development in the adult brain. Besides, LIS1, NDE1, NDEL1 (Brandon et al., 2004; Burdick et al., 2008), PDE4B (Fatemi et al., 2008; Pickard et al., 2007), and Dixdc1 (Singh et al., 2010) are previously reported to be physically interacted with DISC1. But the protein involved in the function of DISC1 on astrogensis remains elusive.

In this study, we focus on the regulation of astrogenesis by DISC1 at the late stage of embryonic brain development and the underlying molecular mechanisms. Using in utero electroporation and in vitro approaches by DISC1 shRNA or expression plamids, we found that DISC1 regulates astrogenesis in the embryo brain. DISC1 depletion suppresses astrogenesis while DISC1 overexpression promotes the process. Moreover, human and mouse DISC1 overexpression rescued astrogenesis defects caused by mouse DISC1-induced defects. A series of experiments deomonstrated that DISC1 phosphorylates downstream MEK and ERK of the RAS signaling and then upregulates the expression of astrocyte related genes. Furthermore, we identified that RASSF7 served as the molecule directly interacting with DISC1. Interaction studies revealed that C terminal domain of DISC1 was directly associated with the C terminal domain of RASSF7, a RAS association protein. In summary, these results represent DISC1 is required for astrogenesis in the embryonic brain and shed light on understanding the pathophysiology of brain diseases.

## RESULTS

### DISC1 Expression in Brain Development

To test whether DISC1 regulates astrogenesis at the late embryonic development stage, we first examine the correlation of DISC1 expression and astrogenesis in brain. Previous studies indicate that DISC1 is crucial to embryogenesis and organ development (Pickard et al., 2007; Singh et al., 2010), we analyzed the DISC1 expression in different tissues and different stages of brain development via western blot analysis. The results showed that DISC1 expression was much higher in the brain than in other tissues (Fig. 1A). DISC1 expression increased from E12 to E15, maintained high level at E17 and P1, and decreased after P7 (Fig. 1B). GFAP expression was increased during this period (E15-P7). These results suggest that the initiation of astrogenesis might be corresponding to the expression of DISC1. Moreover, the *in vivo* immunostaining of brain slices at E16 and P2 showed that DISC1 was highly expressed and restricted in the ventricular zone (VZ)/subventricular zone (SVZ). Furthermore, the colocalization of DISC1 and GFAP was examined in SVZ at P2 (Fig. 1C). These data suggest that DISC1 might be involved in broad brain development, including astrocyte differentiation.

**Figure 1.**
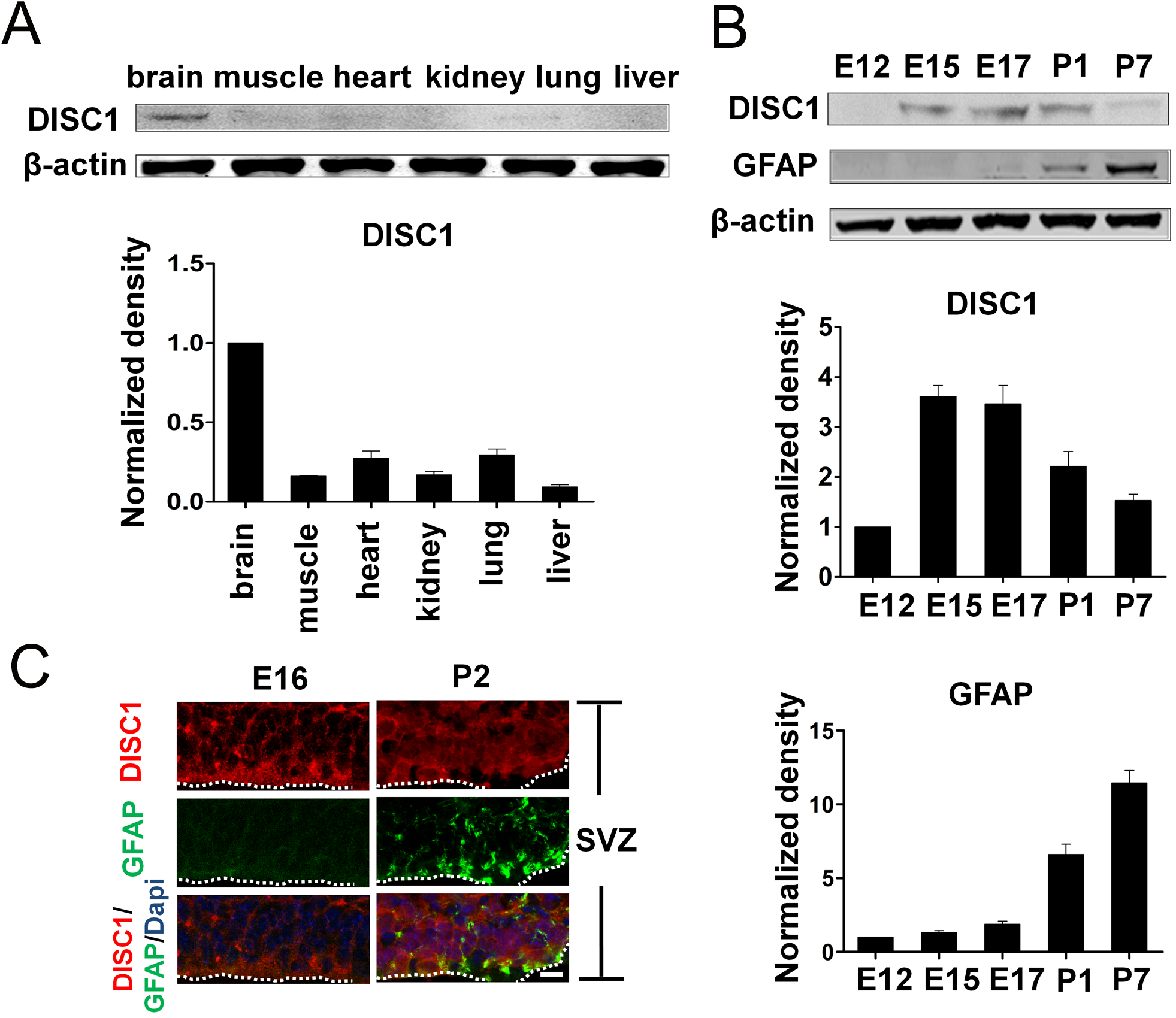
DISC1 expression in brain development. A Western blotting analysis of DISC1 protein in brain, muscle and four other tissues. Quantification of DISC1 expression levels was demonstrated. Error bars indicate s.e.m. (n=3) B Western blot analysis of the level of DISC1 and GFAP protein in the brain cortex from E12 to P7 during embryonic cortical development (E12-P7). Bottom: the graphs demonstrating the quantification of protein expression levels. β-actin was used as an internal control for immunoblotting. Error bars indicate s.e.m. (n=3) C Co-Immunostaining of DISC1 and GFAP in the subventricular zone (SVZ) of brain at E16 or P2. The dotted line indicates the edge of the neocortical SVZ. Scale bar: 50p.m. DISC1, Disrupted in Schizophrenia1; GFAP, glial fibrillary acidic protein; Dapi, 4’,6’-diamidino-2-phenylindole.

### DISC1 Depletion Represses Astrogenesis *in vitro* and *in vivo*

To further determine the regulation of DISC1 on astrogenesis, a series of DISC1 knockdown experiments were performed *in vitro* and *in vivo*. First, four shRNAs were constructed into a lentiviral vector and the knockdown efficiency were examined (Fig. S1A). The result showed that shRNA2 and shRNA3 had high knockdown efficiency. Then, shRNA2 with the highest knockdown efficiency was used for the most of knockdown experiments. We also confirmed that the DISC1 level decreased *in vivo* compared to the control after in utero electroporation (IUE) combined with the DISC1 shRNA2 (Fig. S1B). Next, we studied the effect of DISC1 on astrocyte differentiation after E15 neural precursor cells (NPCs) were infected with the DISC1 shRNA2 lentivirus. Western blotting and GFAP immunocytochemistry showed that DISC1 shRNA2 inhibited GFAP expression, and the number of NPCs that differentiated into astrocytes dramatically decreased with or without LIF treatment (Fig. 2A-B, Fig. S1C). Given that astrogenesis initiates and continues during late embryonic development, control, or DISC1 shRNA2 constructs were electroporated into the lateral ventricle of embryos at E16 *in vivo*. The brain sections were then harvested at P2 and stained for different glial markers of different subtype cells at different stages of astrogenesis. According to previous studies, common progenitor cells give rise to glial progenitor cells, astrocyte progenitor cells, and astrocytes during astrogenesis (Huse and Holland, 2010). Here, we used GLAST, FGFR3, and GFAP to label glial progenitor cells, astrocyte progenitor cells, astrocytes, and mature astrocytes, respectively (Li et al., 2012a; Molofsky et al., 2012). The immunostaining results revealed that the amount of GLAST-positive glial progenitor cells, FGFR3-labeled astrocyte progenitor cells, and GFAP-labeled astrocytes were reduced by 42.4%, 67.8%, and 57.5%, respectively, in response to DISC1 knockdown (Fig. 2C-E). And a concomitant decrease of astrocytes by DISC1 knockdown could be seen in postnatal mouse brains after electroporation at P0, at which neurogenesis is almost finished (Fig. S1D). Moreover, rare morphologically mature astrocytes were observed in brains electroporated with DISC1 shRNA2 (Fig. 2F). To exclude the off target effect, another shRNA3 plasmid was performed to examine astrocyte differentiation in vitro and in vivo. Consist with shRNA2, shRNA3 knockdown decreased GFAP expression and the number of GFAP-GFP positive cells (Fig. S1E-S1F). Collectively, these data suggest that DISC1 is necessary for astrocyte differentiation.

**Figure 2.**
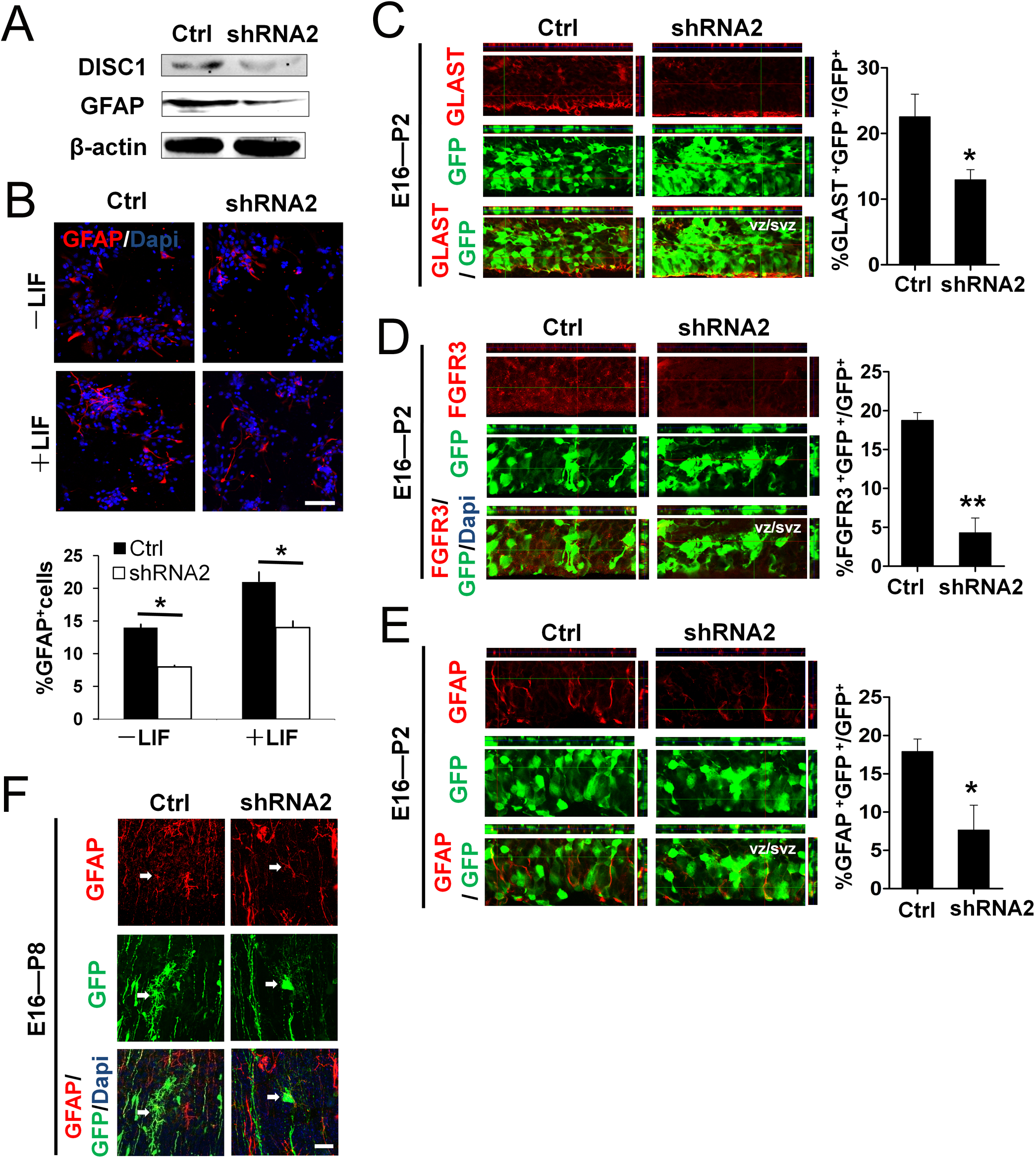
DISC1 knockdown results in astrogenesis defects *in vitro* and *in vivo*. A DISC1 knockdown decreases the levels of DISC1, GFAP in embryonic E15 NPCs infected with lentivirus and cultured in differentiation medium for three days. B DISC1 deficiency induces fewer astrocytes to differentiate from embryonic NPCs treated in glial cell differentiation medium without or with LIF (50 ng/ml) for three days. Top, staining for GFAP displayed astrocyte differentiation. Bottom, the percentages of GFAP-positive astrocytes were measured (*t*-test: *p<0.05). Error bars indicate s.e.m. (n=3). Scale bar: 50p.m. C DISC1 knockdown decreases the number of glial progenitor cells. Left, GLAST immunostaining of P2 cortices electroporated with control or DISC1 knockdown plasmid at E16. Right, the graph showing the proportion of GLAST^+^GFP^+^ glial progenitor cells (*t*-test: *p<0.05). Error bars indicate s.e.m. (n=3). Scale bar: 50μm. D DISC1 knockdown reduces the number of FGFR3-labeled astrocyte progenitor cells. Confocal images of FGFR3 immunostaining are shown on the left, and the percentage of double-labeled FGFR3-GFP cells relative to GFP-positive cells is calculated on the right (*t*-test: **p<0.01). Error bars indicate s.e.m. (n=3). Scale bar: 50μm. E DISC1 knockdown decreases the number of GFAP-labeled astrocytes, as shown in the images of immunostaining. The graph shows percentage of GFAP+GFP+ astrocytes of the GFP^+^ cells (*t*-test: *p<0.05). Error bars indicate s.e.m. (n=4). Scale bar: 50μm. F Images of the astrocyte morphology in P8 brains. The morphology of GFP-labeled mature astrocytes in the brains of mice electroporated with DISC1 shRNA2 plasmid at E16 was not typical comparing with the control. Scale bar: 20μm. LIF, leukemia inhibitor factor; GLAST (Slc1a3), glial high affinity glutamate transporter member 3; FGFR3, fibroblast growth factor receptor 3. See also Supplementary Figure S1.

### DISC1 Overexpression Enhances Astrogenesis

The astrogenesis defect caused by DISC1 knockdown prompted us to perform reciprocal DISC1 gain-of-function experiments. To further analyze the role of DISC1 during astrogenesis, we created a DISC1 overexpression plasmid to efficiently increase the DISC1 expression level (Fig. 3A, Fig. S2A). The results obtained from the immunocytochemistry and western blot analyses showed that DISC1 significantly promoted astrogenesis and increased the expression of GFAP *in vitro* (Fig. 3A-B). The amount of GFAP-positive cells was increased by 48.3% and 55.2% after treatment with and without LIF, respectively. Additionally, DISC1 overexpression *in vivo* by electroporation increased the number of GLAST-positive radial progenitor cells, FGFR3-positive astrocyte precursor cells, GFAP-positive astrocytes (Fig. 3C-E). Moreover, the morphology of astrocytes electroporated with the DISC1 overexpression plasmid was typically more mature and complex *in vivo* (Fig. 3F). Taken together, these results indicate that DISC1 overexpression facilitates astrogenesis both *in vitro* and *in vivo*.

**Figure 3.**
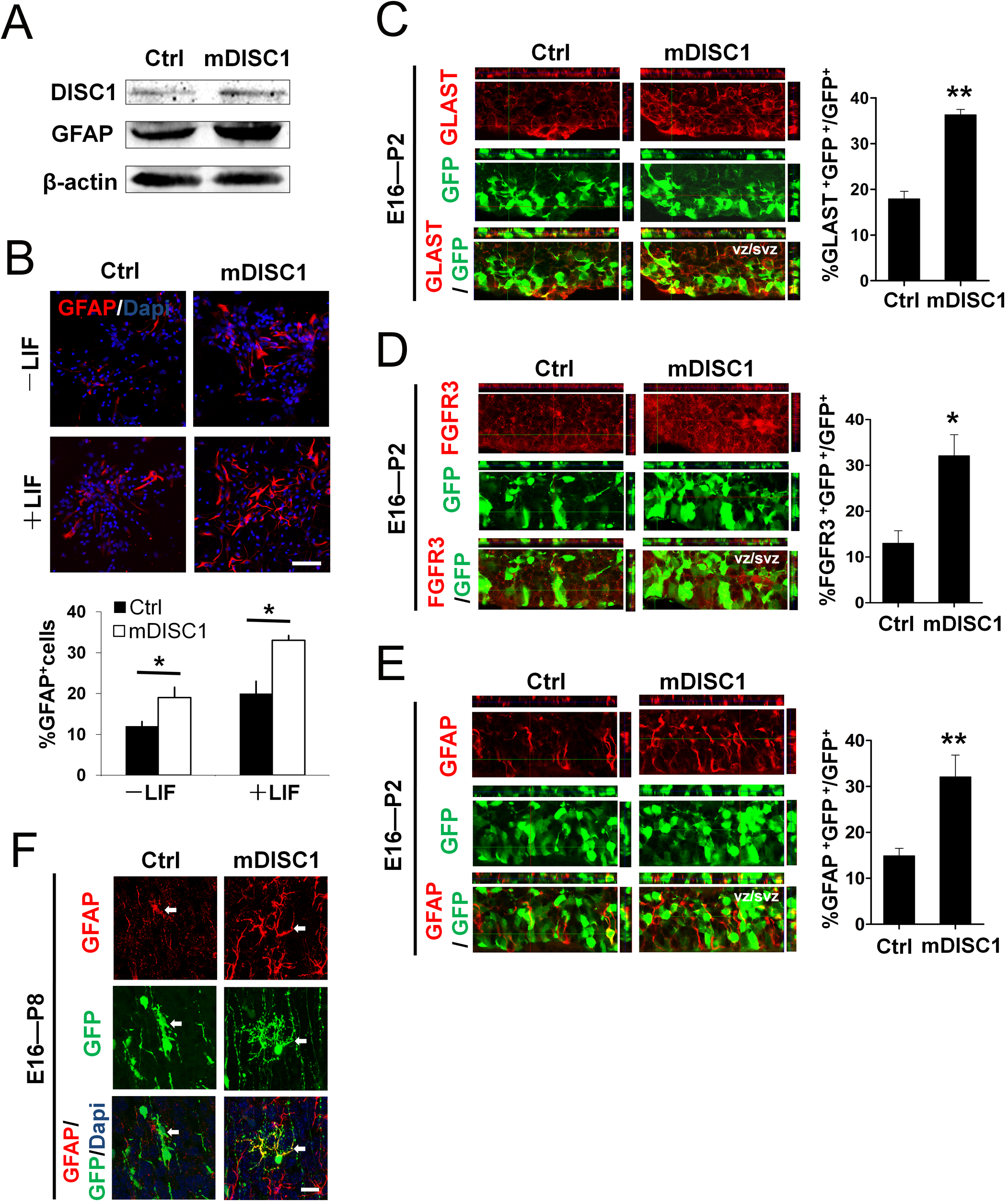
DISC1 overexpression enhances astrogenesis. A E15 NPCs were infected with lentivirus and cultured in differentiation medium for three days. Western blotting showed elevated levels of DISC1, GFAP due to DISC1 overexpression. β-actin was used as an internal control for immunoblotting. B DISC1 overexpression induces more astrocytes differentiation from E15 NPCs treated in differentiation medium without or with LIF (50 ng/ml) for three days. Top, staining for GFAP displayed astrocyte differentiation. Bottom, the percentages of GFAP-positive astrocytes were measured (*t*-test: *p<0.05). Error bars indicate s.e.m. (n=3). Scale bar: 50μm. C-E DISC1 overexpression enhances astrogenesis *in vivo*. E16 embryonic brains electroporated with control and DISC1 overexpression plasmids were analyzed at P2. The left images in (C-E) were immunostaining for GLAST, FGFR3, GFAP, and respectively. The proportion of GLAST^+^GFP^+^ glial progenitor cells, FGFR3^+^GFP^+^ cells, and GFAP^+^GFP^+^ astrocytes to GFP^+^cells is shown in the right graphs in (C-E), respectively (*t*-test: *p<0.05, **p<0.01). Error bars indicate s.e.m. (n=3). Scale bar: 50μm. F Images of astrocyte morphology in P8 brains. The morphology of GFP-labeled astrocyte in the brains electroporated with DISC1 overexpression plasmid at E16 was more mature than that in the control brain. Scale bar: 20μm. See also Supplementary Figure S2.

### DISC1 Overexpression Rescues Astrogenesis Defects Caused by DISC1 Depletion

To further examine DISC1 as a key regulator in astrogenesis, we generated a human DISC1 (hDISC1) overexpression plasmid and performed rescue experiments. First, western blotting and immunostaining were used to examine the effect of hDISC1 on astrogenesis. The results showed a higher level of GFAP expression and an increased number of GFAP-labeled astrocytes when hDISC1 was overexpressed (Fig. 4A-B). After IUE *in vivo*, we observed that GFAP expression and the astrocyte number were increased when hDISC1 and DISC1 shRNA were co-expressed (Fig. 4C-D). Additionally, mDISC1 (mouse DISC) could also rescue the astrocyte differentiation defect caused by DISC1 knockdown (Fig. S2B). These data suggest that mDISC1 or hDISC1 overexpression could restore the astrogenesis defects caused by mDISC1 knockdown.

**Figure 4.**
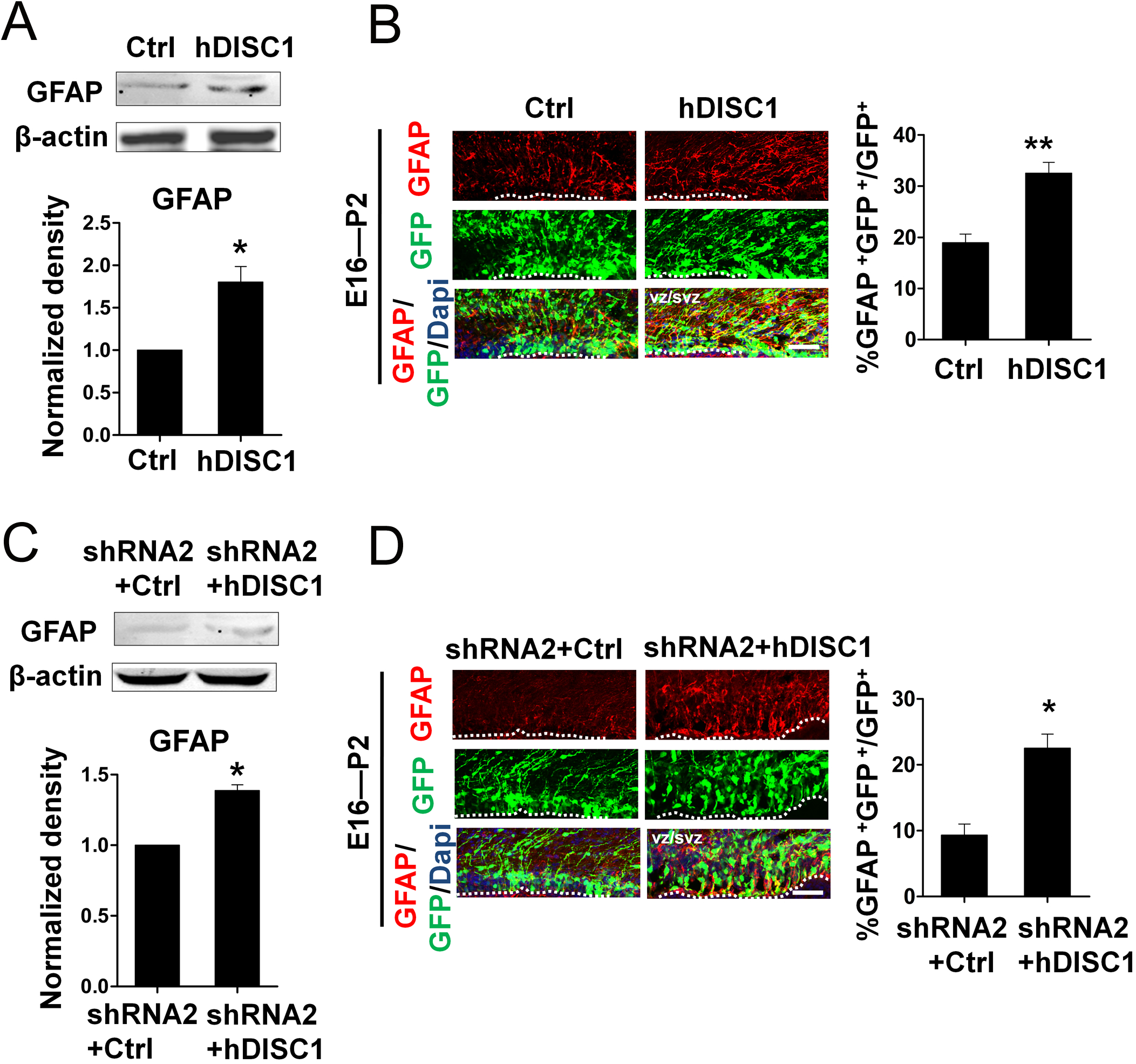
DISC1 overexpression rescues astrogenesis defects induced by mouse DISC1 depletion. A,C Reduced GFAP expression due to mouse DISC1 (mDISC1) knockdown was rescued by human DISC1 (hDISC1) overexpression. Embryonic NPCs were infected with different combinations of lentivirus and cultured for four days. The GFAP protein expression was quantified (*t*-test: *p<0.05). Error bars indicate s.e.m. (n=3). B The overexpression of hDISC1 leads to more astrocytes *in vivo*. Left, increased numbers of GFAP-positive cells in embryo cortices electroporated at E16. Right, the proportion of GFAP^+^GFP^+^ astrocytes (*t*-test: **p<0.01). Error bars indicate s.e.m. (n=4). Scale bar: 50μm. D The decrease in astrocyte number caused by mDISC1 knockdwon is rescued by a gain of hDISC1 *in vivo*. Left, confocal images of GFAP immunostaining. Right, the percentage of double-labeled GFAP-GFP cells relative to GFP cells (*t*-test: *p<0.05). Error bars indicate s.e.m. (n=3). Scale bar: 50μm.

### DISC1 Regulates Astrogenesis by Modulation of pMEK and pERK Levels

Multiple signaling pathways are reportedly involved in gliogenesis, including the JAK/STAT, MEK/ERK, and Notch pathways (Bonni et al., 1997; Li et al., 2012b; Zhou et al., 2010). Li *et al*. found that the RAS/MEK/ERK signaling pathway plays a crucial role in activating the expression of astrocyte-specific genes (Li et al., 2012b). Furthermore, the activation of RAS, MEK, and ERK are three major nodes in the pathway. To test whether DISC1 regulates astrogenesis via this pathway, we checked the level of phosphorylated and total ERK after treatment with LIF. The data showed that LIF activated ERK (Fig. S3A). Western blot analysis showed that the expression of pMEK and pERK decreased due to DISC1 knockdown and increased due to DISC1 overexpression (Fig. 5A-B, Fig. S3B). We further observed that the overexpression of ERK could rescue the deficits of astrocytes caused by DISC knockdown *in vivo* (Fig. 5C). The active form of ERK (pERK) is reported to function following translocation into the nucleus (Clark et al., 2004). Thus, we detected the location of pERK treated with LIF for different time intervals after DISC1 was knocked down or overexpressed. The results showed that pERK was primarily located in the cytoplasm in the absence of treatment, transported to the nucleus after 10min of short-term treatment with LIF, and increasingly re-localized to the cytoplasm after 2 h (Fig. S3C). Collectively, these results indicate that DISC1 regulates astrogenesis by modulation of the pMEK and pERK levels.

**Figure 5.**
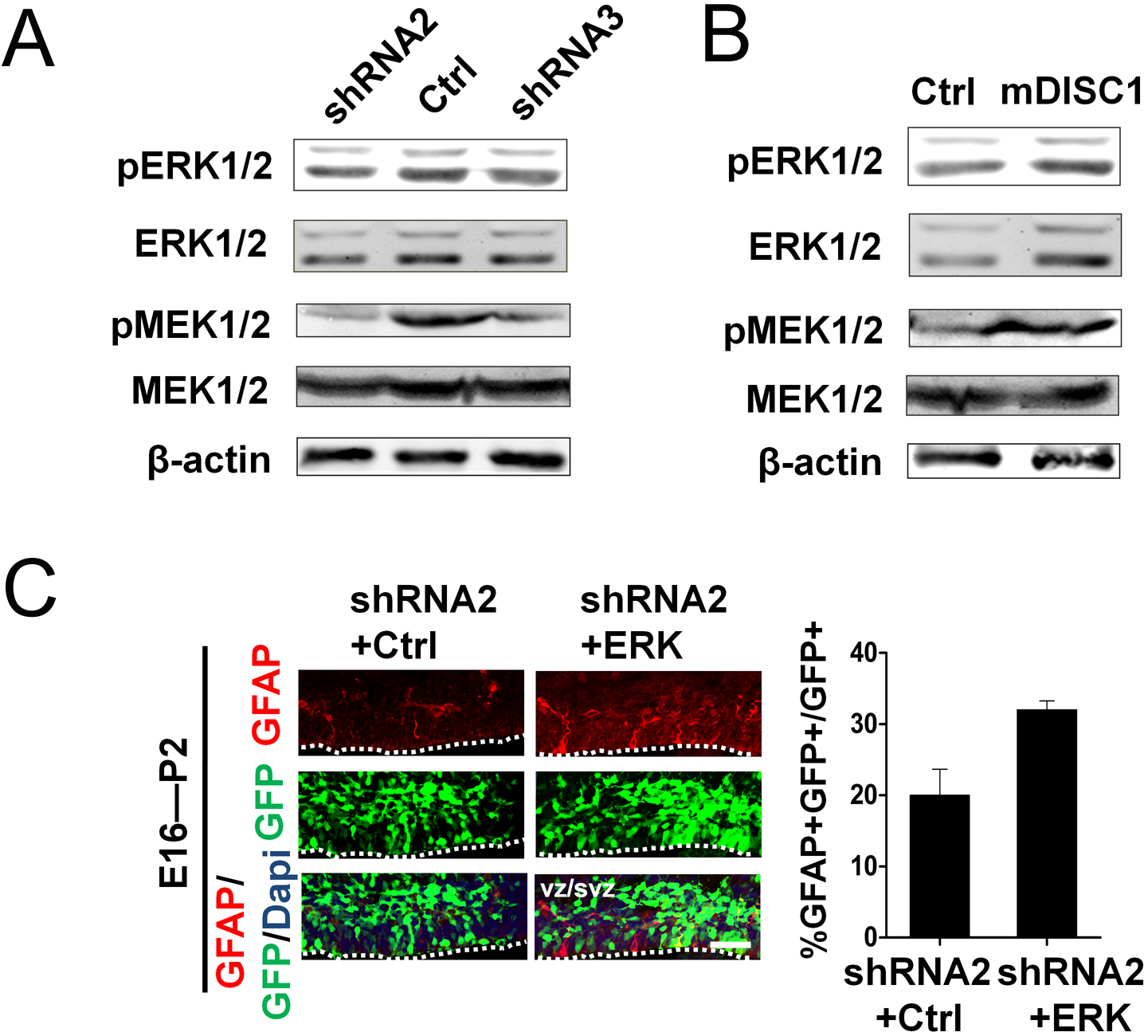
DISC1 regulates astrogenesis by modulation of pMEK and pERK levels. A Western blot analysis of MEK and ERK. Left, NPCs infected with DISC1 shRNA2 or shRNA3 lentivirus and cultured in differentiation medium for three days. Following by serum starvation overnight, cells were treated with LIF (50 ng/μl) for 10min. The cell lysates were subjected to western blot analysis for phosphorylated and total MEK, and phosphorylated and total ERK. The levels of pMEK and pERK decreased due to DISC1 knockdown. B Western blotting of NPCs infected with the DISC1 overexpression lentivirus showed increased levels of MEK and ERK phosphorylation. C The decrease in astrocytes in response to DISC1 shRNA2 was rescued by ERK overexpression *in vivo*, as demonstrated by GFAP staining. Right, the percentages of GFAP^+^ GFP^+^ cells relative to GFP^+^ cell numbers (*t*-test: *p<0.05). Error bars indicate s.e.m. (n=3). Scale bar: 50 μm. See also Supplementary Figure S3.

### DISC1 Regulates RAS/MEK/ERK Signaling Pathway by Direct Interaction with RASSF7

Co-IP analysis showed that DISC1 was not directly interacted with ERK, MEK or c-RAF (Fig. S3D). Therefore, we hypothesize that DISC1 may directly interact with an upstream component of the RAS pathway. Human RASSF7, a member of the RAS-association domain family, was previously reported as a potential binding partner of human DISC1 based on yeast two-hybrid screens (Morris et al., 2003). Furthermore, the similarities of human DISC1 and mouse DISC1 as well as human RASSF7 and mouse RASSF7 are 54.76% and 73.08%, respectively. First, we investigated whether there was direct interaction between RASSF7 and RAS and the effect of RASSF7 on the phosphorylation of MEK and ERK. Our co-IP analysis results showed that RASSF7 was indeed interacted with RAS (Fig. 6A). We next investigated whether there was an interaction between DISC1 and RASSF7 and their involvement in astrogenesis. To address this possibility, we generated recombinant Flag-tagged DISC1, HA-tagged DISC1, Flag-tagged RASSF7, and HA-tagged RASSF7. The results of co-immunoprecipitation (co-IP) showed that DISC1 and RASSF7 directly interacted with each other (Fig. 6B-C). Moreover, the co-localization of DISC1 and RASSF7 was observed in cells co-transfected with Flag-tagged DISC1 and HA-tagged RASSF7 (Fig. S4A). And the binding complex of DISC1-RASSF7 gradually increased as the levels of RASSF7 increased (Fig. S4B). Furthermore, the result of in utero electroporation demonstrated that RASSF7 overexpression enhanced astrocyte differentiation *in vivo* (Fig. 6D).

**Figure 6.**
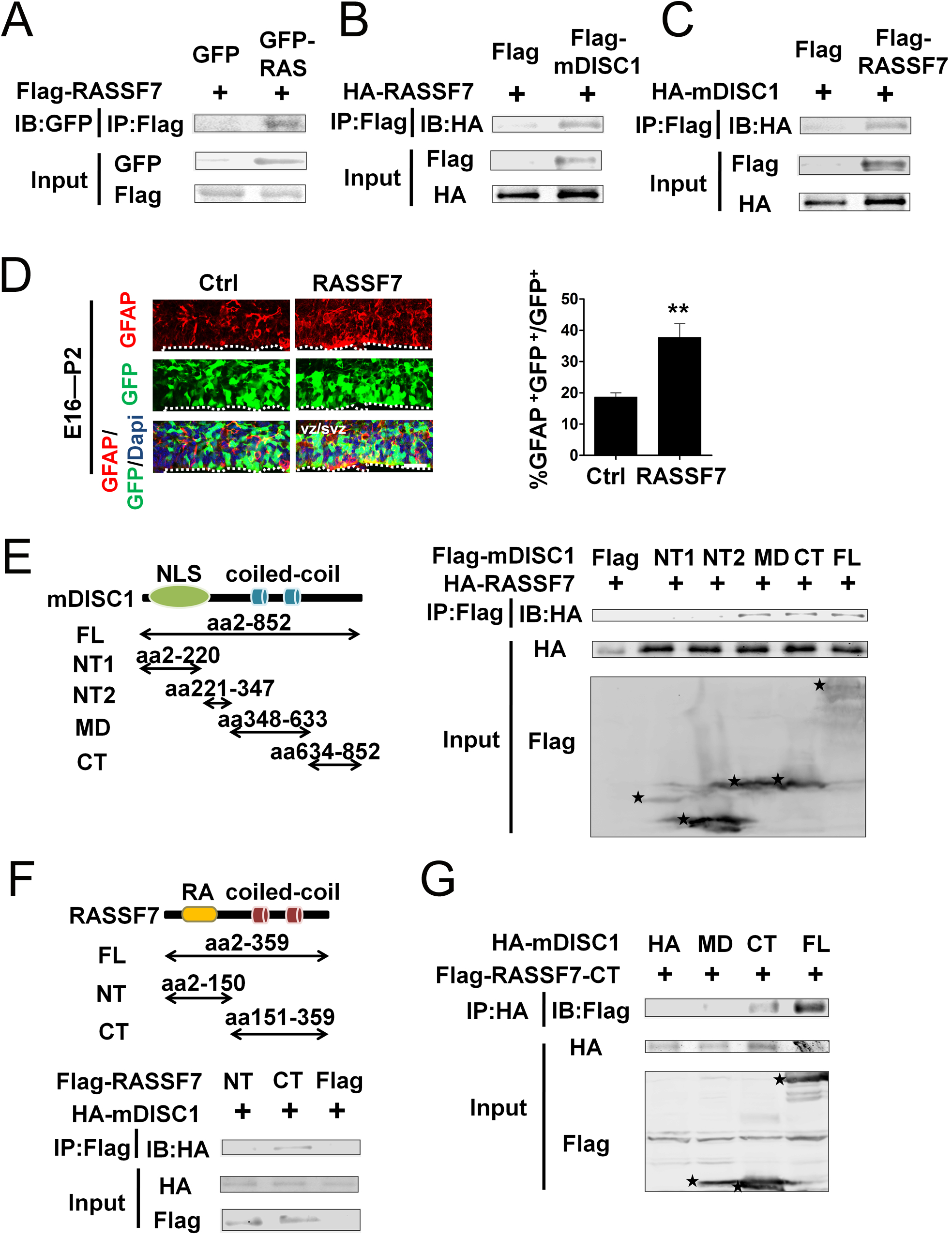
DISC1 regulates RAS/MEK/ERK signaling pathway by direct interaction with RASSF7. A Direct interaction between RAS and RASSF7. Lysates from NPCs infected with Flag-RASSF7 and GFP-vector or GFP-RAS lentivirus and differentiated for three days were immunoprecipitated with a GFP antibody and immunoblotted with a Flag antibody. RASSF, the RAS-association domain family. B-C Direct interaction between DISC1 and RASSF7. B, NPCs infected with full-length HA-tagged RASSF7 lentivirus together with full-length Flag-tagged DISC1 lentivirus or Flag-vector lentivirus were cultured in differentiation medium for three days. Cell lysates were subjected to co-IP using antibodies against Flag and immunoblotted for HA-RASSF7. C, co-IP of lysates from cells infected with full-length HA-tagged mDISC1 lentivirus together with full-length Flag-tagged RASSF7 lentivirus or Flag-vector lentivirus. D RASSF7 overexpression increases the number of astrocytes *in vivo*. The left images show GFAP-labeled astrocytes and the right graph shows the percentage of GFAP^+^GFP^+^ astrocytes relative to the number of GFP+cells (*t*-test: **p<0.01). Error bars indicate s.e.m. (n=3). Scale bar: 50 μm. E Left, A schematic diagram of the Flag-tagged DISC1 recombinant fragments used for domain mapping. Right, RASSF7 binds to the CT region of DISC1 between residues 634 and 852. Lysates from NPCs co-infected with HA-RASSF7 lentivirus and each of the Flag-tagged DISC1 domains were immunoprecipitated with an antibody against Flag and immunoblotted with an antibody against HA. F Top, A schematic showing the different domains and regions of RASSF7 used to generate Flag-tagged fragments. Bottom, DISC1 preferentially binds to the C-terminus of RASSF7 (CT region). A western blot of the *in vitro* co-IP of recombinant HA-tagged DISC1 with different recombinant Flag-tagged fragments. G Direct interaction between the fragments of DISC1 and RASSF7. Lysates from NPCs cells co-infected with the Flag-tagged RASSF7-CT fragment lentivirus together with full-length HA-tagged DISC1 lentivirus, the HA-tagged DISC1-CT fragment lentivirus,or Flag-vector lentivirus were subjected to co-IP using antibodies against HA and immunoblotted for Flag-RASSF7-CT. See also Supplementary Figure S4.

**Figure 7.**
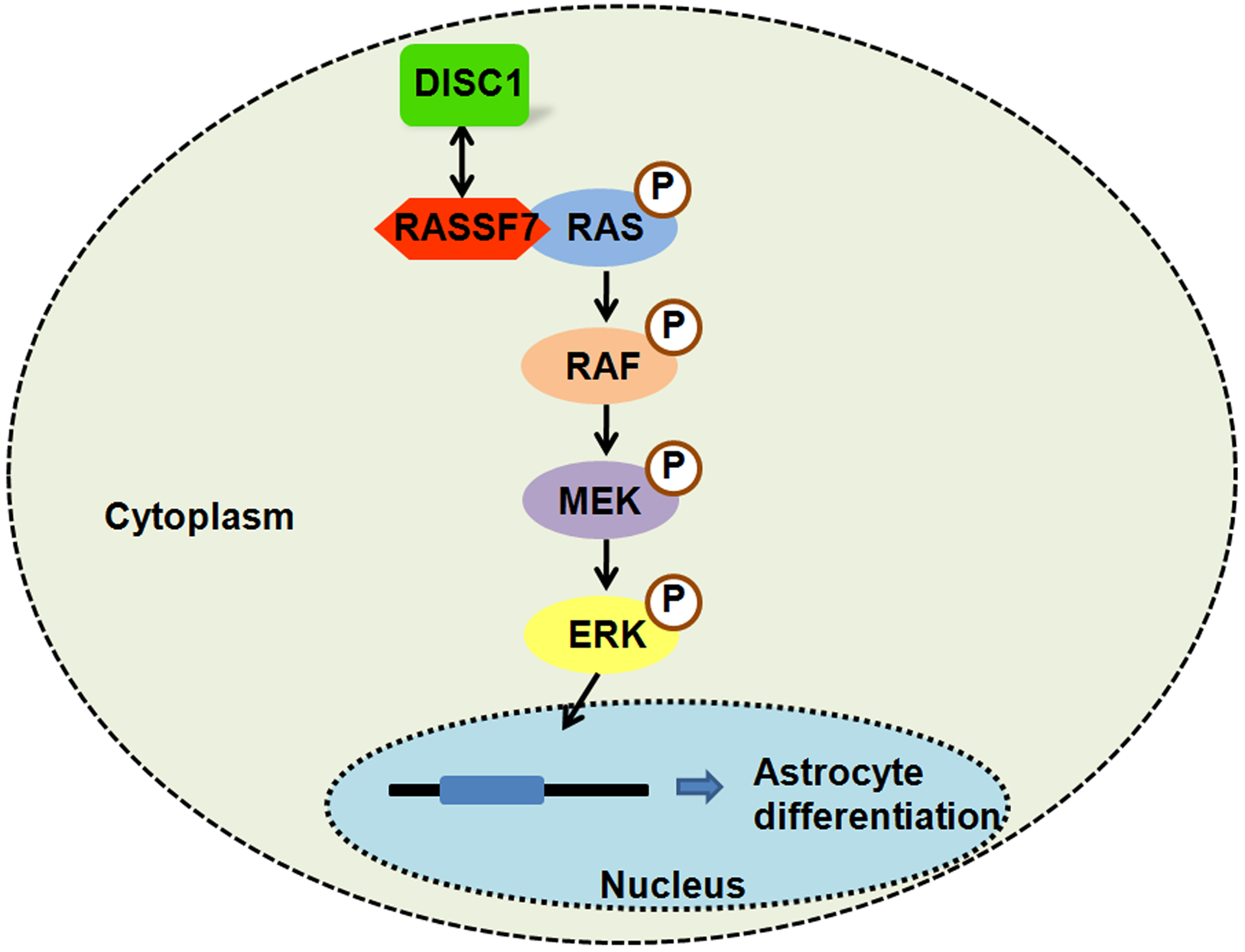
A model of DISC1 function in astrogenesis during cortical development.

How does DISC1 interact with RASSF7? DISC1 contains multiple interaction domains: a globular domain localized in the N-terminus, two Leucine zipper domains localized in the middle and C-terminus, and several coiled-coil domains along the full-length (Singh et al., 2010). To map the region(s) of DISC1 involved in the association with RASSF7, we generated a series of Flag-tagged fragments of DISC1 and co-expressed them with HA-tagged RASSF7. The co-IP results revealed that the CT domain of DISC1 was associated with RASSF7 in the protein complex. In addition, the MD domain of DISC1 also directly bound to RASSF7 (Fig. 6E). RASSF7, originally named HRC1, contains a RAS-association domain located at its N-terminus (Sherwood et al., 2008). To map the RASSF7 domain(s) involved in RASSF7-DISC1 association, we also generated a series of Flag-tagged fragments of RASSF7 and co-expressed them with HA-tagged DISC1 in progenitor cells. The co-IP results demonstrated that the CT domain of RASSF7 was involved in the association with DISC1 in the complex (Fig. 6F). Furthermore, the co-IP results demonstrated that the CT domain of DISC1 was sufficient to associate with the CT domain of RASSF7 (Fig. 6G). Following, we constructed DISC1-ΔNT2 or DISC1-ΔMD plasmids and performed rescue experiments to test whether DISC1 mutants could rescue the mDISC1 shRNA effects in vivo. Immunostaining of GFAP showed that the astrocyte number was increased when DISC1-**Δ** NT2 and mDISC1 shRNA2 were co-expressed (Fig. S4C). However, the number of GFAP+ GFP+ cells was not significantly affected when DISC1-ΔMD and mDISC1 shRNA2 were co-expressed (Fig. S4D). These data suggested that MD domain, not NT2 domain, was important for the function of DISC1. Taken together, these results indicate a direct and complex interaction between DISC1 and RASSF7 via multiple domains.

Since RASS7 belongs to RAS family, we then ask whether RAS/MEK/ERK signaling is involved in the process. First, we examined pERK and found that the levels of pERK were increased in the presence of RASS7 with a dose dependent manner (Fig. S4E). Moreover, the results showed that the overexpression of RASSF7 or DISC1 alone increased the levels of pMEK and pERK in the presence of LIF. And RASSF7 and DISC1 overexpression results in higher levels of pMEK and pERK (Fig. S4F). Interestingly, the overexpression of RASSF7 restored the pMEK and pERK levels caused by the reduction of DISC1 shRNA (Fig. S4G). The data suggest that RASSF7 is required for the DISC1-dependent modulation of MEK/ERK signaling.

## DISCUSSION

*DISC1* is one of the few single-genes definitively associated with psychiatric disease, such as schizophrenia, bipolar, and unipolar depression (Bradshaw and Porteous, 2012). Previous studies report that mutant DISC1 plays a dominant negative role in inducing a DISC1 loss of function in schizophrenia (Kamiya et al., 2005). However, it is unclear how DISC1 dysfunction results in a series of mental disorders. DISC1 has been demonstrated to widely function in neuronal development, but its role in astrogenesis is not clear. Furthermore, the etiology of schizophrenia remains to be comprehensively characterized. In this study, we show that DISC1 plays an important role in astrocyte progenitor differentiation during late embryonic brain development. Suppression of DISC1 expression inhibits astrogenesis *in vitro* and *in vivo*, whereas DISC1 overexpression substantially enhances the process. Moreover, mouse and human DISC1 overexpression rescued astrogenesis defects caused by DISC1 knockdown in mice.

In addition, we also checked neural stem cell proliferation and neuron differentiation. E16 embryonic brains electroporated with control and DISC1 knockdown plasmids were analyzed at E18 or P0 (Fig. S5A-B). The results showed the cells mainly located at the VZ/SVZ and a small part of cells migrate to the IZ. We found the number of Nestin-positive cells was slightly decreased when DISC1 was knocked down or slightly increased when DISC1 was overexpressed when brains were electroporated at P0 and analyzed at P3 (Fig. S6A-B). The data suggest that DISC1 knockdown does not affect Nestin positive progenitor cell proliferation at late stage. We also examined the neuron differentiation at late stage. The results showed that the number of neurons differentiated from isolated P0 NPCs was not significantly affected in vitro by DISC1 knockdown or overexpression (Fig. S6C-F).

Gliogenesis occurs during the late stages of embryogenesis after neurogenesis is nearly finished, when NPCs generate more astrocytes and fewer neurons. The specific genes related to astrocyte development were temporally and spatially expressed during late embryonic brain development following the activation of different relevant intrinsic signaling pathways. Among these pathways, JAK-STAT pathway is important for controlling the onset of astrogenesis (Fan et al., 2005). In addition, the RAS/MEK/ERK signaling pathway has been identified to regulate gliogenesis (Li et al., 2012b), but the underlying molecular mechanism remains to be further explored. Moreover, many uncertainties remained in the intrinsic factors that change the RAS/MEK/ERK signaling pathway. Our data indicate that DISC1 regulates astrogenesis by modulating the pMEK and pERK levels. Furthermore, consistent with a previous study (Clark et al., 2004), we confirmed that the active form of ERK begins to function following translocation into the nucleus. Interestingly, DISC1 is participated in this pathway by interation with RASSF7, a member of the RAS-association domain family. A previous study used yeast two-hybrid screening to identify that hRASSF7 interacts with hDISC1. We compared the similarities of protein sequences and found that human DISC1 and mouse DISC1 as well as human and mouse RASSF7, were highly homologous. Based on this information, co-immunostaining was performed and displayed that mouse DISC1 perfectly interacts with RASSF7.

To date, lines of evidence have provided multiple signaling pathways that correlated with the intrinsic pathogenesis of schizophrenia (Kyosseva, 2004). Recently, DISC1 was proven to be involved in GSK3β/β-Catenin signaling (Mao et al., 2009a), AKT-mTOR signaling (Kim et al., 2009), and GABA signaling. Given the variety of proteins that bind DISC1, it plays many roles in the central nervous system via these signaling pathways. Dysfunction in these signaling pathways resulting from mutant DISC1 contributes to the development of schizophrenia. Here, we add that DISC1 regulates RAS/MEK/ERK pathway, which is important for brain astrocyte development and function. The influence of DISC1 on the development of neurons and oligodendrocytes has been investigated (Hattori et al., 2014; Mao et al., 2009a). Here, we further discover the effects of DISC1 on astrocyte development. In summary, our results provide a framework for understanding the effect of DISC1 on astrogenesis may contribute to brain development and the etiology of psychiatric disorders.

## MATERIALS AND METHODS

Detailed information can be found in Supplementary Materials and Methods.

### Mice

ICR female mice were used for in utero and postnatal electroporation experiments. All animal experiments and protocols were approved by the Animal Care and Use Committee of Institute of Zoology, Chinese Academy of Sciences.

### In utero and postnatal electroporation

Every embryo was electroporated with five 50 ms pulses at 50V with 950 intervals, using 5 mm paddle electrodes. This manipulation was finished in 30 min before the embryos were returned to the abdominal cavity (Wang et al., 2014). For neonatal pups, they were electroporated with four 50 ms pulses at 90V with a 950 ms interval.

## SUPPLEMENTAl INFORMATION

Supplemental information for this article is available online:

## ACKNOWLEDGEMENTS

We are grateful to Shiwen Li for help with confocal microscopy. This work was supported by the Ministry of Science and Technology of China (2015CB964501 and 2014CB964903), the National Science Foundation of China (31371477), and the Strategic Priority Research Program (XDA01020301).

## AUTHOR CONTRIBUTIONS

SW, QL, and JJ designed the study, analyzed the date, wrote the paper; SW, QL, HQ, and TS performed the research; HL, and FJ analyzed parts of the data.

